# Low-intensity environmental education can enhance perceptions of culturally taboo wildlife

**DOI:** 10.1101/757724

**Authors:** Samual T. Williams, Kathryn S. Williams, Natasha Constant, Lourens H. Swanepoel, Peter J. Taylor, Steven R. Belmain, Steven W. Evans

**Author notes:** Corresponding authors: Samual Williams & Lourens Swanepoel.

## Abstract

Traditional cultural beliefs influence perceptions of animals, and in some cases can result in persecution of wildlife. Stigmas against species associated with witchcraft, for example, can act as a barrier to the uptake of more cost effective, sustainable, and environmentally sound practices such as reducing crop damage by controlling rodent agricultural pests by relying on indigenous predators rather than pesticides. One way of enhancing perceptions of wildlife to increase participation in such ecologically-based rodent management (EBRM) schemes, is the development of environmental education initiatives. Low intensity programmes are cost-effective and can produce positive attitudinal shifts, but their impact has not been assessed for species strongly associated with witchcraft. We set out to test whether a single presentation on the natural history of owls (order Strigiformes) could improve perceptions of these species, and increase willingness to participate in an EBRM scheme that involved the installation of owl boxes to increase owl populations and reduce rodent populations and crop damage in agricultural fields. We used a questionnaire survey to assess perceptions of owls at four schools in two villages in South Africa. Our initial survey sampled perceptions of respondents before listening to the presentation. A follow-up survey conducted three months later sampled the perceptions of respondents that had listened to the presentation as well as perceptions of a control group that did not listen to the presentation. We found that associations between owls and witchcraft was a common theme driving negative perceptions of owls. Respondents that watched the presentation had more positive perceptions of owls than respondents that had not watched the presentation, and they were more likely to be willing to put up owl boxes near their home. Despite this shift, negative perceptions of owls still dominated responses due to cultural associations with the occult. These findings indicate that even low-intensity programmes can be effective at enhancing perceptions of wildlife associated with witchcraft. We suggest that environmental education programmes featuring culturally taboo species should adopt a culturally sensitive and locally tailored approach, focus on the benefits these species provide, and may be more effective when delivered with greater intensity.

## 2. Introduction

Human interactions with the natural world are frequently rooted in rich cultural interpretations (Becken *et al*., 2013; Garibaldi and Turner, 2004; Risiro *et al*., 2013). Historically, cultural interpretations of nature were overlooked in conservation strategies by pursuing an exclusionary ‘fortress’ approach to environmental management and reducing cultural constructs to non-tangible ideologies (Binnema and Niemi, 2006; Drew and Henne, 2006; Tyrrell, 2010). However, cultural interpretations of animals can have significant implications for wildlife conservation, especially when they manifest in physical outcomes that either protect or endanger species (Dickman, 2010; Madden and McQuinn, 2014). Even cultural interpretations of imaginary creatures can impact wildlife conservation (Holmes *et al*., 2018). Recent research highlights the futility of pursuing a purely biological based conservation agenda without addressing relevant cultural considerations, especially in situations of human-wildlife conflict (Dickman, 2010; Madden and McQuinn, 2014; Peterson *et al*., 2010). Consequently, conservationists are beginning to recognise the need to integrate social science perspectives into their work and adopt a more interdisciplinary approach (Balmford and Cowling, 2006; Mascia *et al*., 2003; White and Ward, 2011), although uptake can be slow (Montgomery *et al*., 2018).

Cultural beliefs such as those linked to witchcraft and the occult are integral to many cultures, especially in Africa, and although mythologies vary substantially, associations between animals and the supernatural are common (Dickman *et al*., 2013; Geschiere, 1997; Kesby, 2003). In southern African societies uneven access to material possessions, wealth, education, and basic healthcare persist, partially as legacies of colonialism and apartheid (Gradin, 2014; Özler, 2007; Rust *et al*., 2016). Unequal distribution of resources creates fears and jealousies which may be expressed through accusations of witchcraft or soul eating, negatively eroding social structures within communities (Geschiere, 1997; Schmoll, 1993; Smith & Andindilile, 2017). Witches are defined as human beings consumed by jealousy, greed, malice, and antisocial tendencies who use supernatural powers to harm others (Ashforth, 2005; Hickel, 2014; Niehaus *et al*., 2001). Witches are excluded from personhood due to their immoral and antisocial characteristics, and their ability to transform into animal familiars (Niehaus *et al*., 2001).

Witches use animals in multiple ways: animals may signify a witch’s presence; animals can act as a witch’s familiar; animal parts are used in *muti* (traditional medicine); and after death, animals may be become a vessel for the witch’s spirit to spread further malevolence (Morris, 2000; Niehaus *et al*., 2001). Predators, nocturnal species, and animals considered dangerous such as owls, hyaenas (family Hyaenidae), cats (family Felidae), or snakes (suborder Serpentes) are most commonly associated with the occult (Cumes, 2004; Niehaus *et al*., 2001). Under some circumstances, fear of these animals is more strongly associated with the human construct of witchcraft than the animals’ biological characteristics (Williams, 2017). Animals associated with witches are often killed as a precautionary measure or for use in traditional medicine (Mikkola and Mikkola, 1997; Williams *et al*., 2013).

Ironically, some of the species most heavily persecuted due to associations with witchcraft provide valuable services to rural communities such as controlling agricultural pests (Muñoz-Pedreros *et al*., 2018; Williams *et al*., 2018b). Smallholder agriculture supports the majority of impoverished people in rural areas (Tscharntke *et al*., 2012; World Bank, 2007), but one of the key constraints on food production to smallholder farmers, is crop damage caused by pests such as rodents (Swanepoel *et al*., 2017). Existing pest control in these areas tends to rely heavily on rodenticides, but such practices cause environmental contamination, poisoning of non-target species, and result in the development of physiological and behavioural resistance to the products used (Buckle and Smith, 2015). To overcome these problems, an alternative approach termed ecologically-based rodent management (EBRM) was developed (Singleton *et al*., 1999), which emphasises more sustainable pest management solutions such as biological control by native species such as mammalian carnivores (order Carnivora) (Williams *et al*., 2018b) or avian predators such as owls (order Strigiformes) (Labuschagne *et al*., 2016). The density of rodents near crop fields as well as crop damage can be reduced, for example, by erecting artificial nest boxes for owls (Labuschagne *et al*., 2016; Paz *et al*., 2013). The adoption of EBRM strategies for rodent pest management can therefore effectively reduce rodent damage whilst decreasing reliance on rodenticides (Jacob *et al*., 2010; Taylor *et al*., 2012).

Superstitious cultural beliefs about species can act as a barrier to the acceptance of using the pest control ecosystem services offered by these species (Williams *et al*., 2018b). One tool that can be used to enhance perceptions of wildlife is environmental education, which can increase knowledge and provoke positive behavioural changes in recipients and their families (Boudet *et al*., 2016; Lawson *et al*., 2019; Rakotomamonjy *et al*., 2015). Education schemes have been successfully applied to increase rates of EBRM adoption; boosting crop yields as a result (Flor and Singleton, 2011). However, environmental knowledge does not always equate to positive conservation actions (Knapp and Poff, 2001). Farmers in Kenya who demonstrated greater knowledge about owl diets were more likely to use pesticides and kill owl prey than farmers with lower levels of understanding (Ogada and Kibuthu, 2008). This trend was attributed to a lack of knowledge about the interrelationships in ecological processes (Ogada and Kibuthu, 2008). Improving EBRM uptake is reliant on delivering education that stresses the benefits native species can provide farmers and emphasises the ecological interconnectedness between rodents, pesticides, and predators (Makundi and Massawe, 2011; Ogada and Kibuthu, 2008).

Long-term education initiatives are expensive, with lack of funding being the major constraint on many programmes (McDuff and Jacobson, 2001), and there are increasing calls to ensure that conservation strategies such as environmental education programmes are cost effective (Cook *et al*., 2017; Naidoo *et al*., 2006). This can be achieved by tailoring environmental education programmes to the audience, available resources, and time constraints (Offord-Woolley *et al*., 2016). Although carefully crafted short-term environmental education programmes can successfully increase knowledge and foster positive environmental perceptions (Farmer *et al*., 2007; Leeds *et al*., 2017; Rakotomamonjy *et al*., 2015), it is yet to be determined if low intensity environmental education can sensitively and cost effectively enhance perspectives of culturally taboo species. Utilising a case study approach, we examined young people’s perceptions of owls in two rural South African communities before and after conducting a low intensity environmental education programme. We assessed whether listening to a single presentation on the natural history of owls, their biological benefits, and the roles of owls and humans within the ecosystem could improve perceptions of these species, and increase willingness to take part in a future EBRM trial to help reduce agricultural damage from rodent pests by installing owl nest boxes. We hypothesise that attitudes towards owls will show a moderate improvement in response to the environmental education scheme, but improvements may be limited by the low intensity of the programme and how deeply entrenched such cultural beliefs tend to be.

## 3. Methods

### 3.1 Study site

We assessed the influence of a low-intensity environmental education programme on perceptions of owls in Ka-Ndengeza (S23.31003 E30.40981) and Vyeboom (S23.15174 E30.39278), two rural villages in Limpopo Province, South Africa (Fig. 1). Vyeboom, located in Makhado municipality, has a population of approximately 5,000, and the most commonly spoken language is Tshivenda (Statistics South Africa, 2019b). In Ka-Ndengeza, Mopani municipality, the population of around 3,500 predominantly speak Xitsonga (Statistics South Africa, 2019a). A small percentage of residents of both villages practice small-scale agriculture, with crops including maize *Zea mays*, peanuts *Arachis hypogaea*, beans *Phaseolus vulgaris*, avocados *Persea americana*, pumpkins *Cucurbita* spp, mangoes *Mangifera* spp, bananas *Musa* spp, litchis *Litchi chinensis*, and oranges *Citrus* spp. Livestock such as cattle *Bos taurus*, donkeys *Equus africanus asinus*, sheep *Ovis aries*, goats *Capra aegagrus hircus*, and poultry such as chickens *Gallus gallus domesticus* were also kept (Williams *et al*., 2018b).

**Fig. 1.**
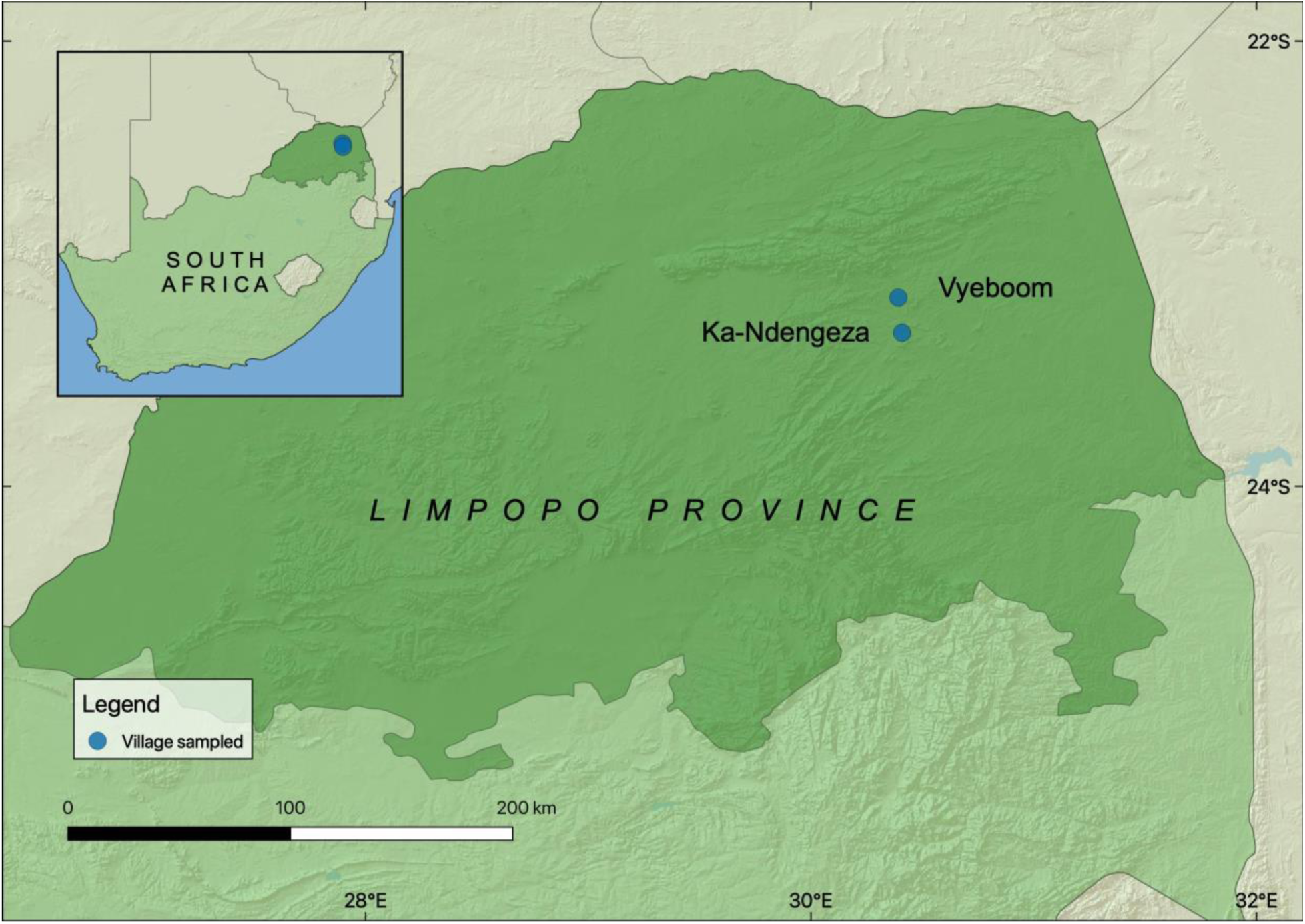
Map of the study sites showing the location of the villages included in the study.

Rainfall at both sites is approximately 700–800 mm per year. There is a hot wet season from October to March and a cool dry season from May to August (Hijmans *et al*., 2005). The main vegetation types are Granite Lowveld and Gravelotte rocky bushveld (Mucina and Rutherford, 2006). Natural vegetation in uplands is dominated by *Combretum zeyheri* and *C. apiculatum* woodlands, while low lying areas are characterised by dense thicket to open savanna dominated by *Senegalia* (*Acacia*) *nigrescens*, *Dichrostachys cinerea*, and *Grewia bicolor* (Mucina and Rutherford, 2006; Williams *et al*., 2018a). Of the 12 species of owls that occur in South Africa, four are found at the study sites: the western barn owl *Tyto alba*, spotted eagle-owl *Bubo africanus*, Verreaux’s eagle-owl *Bubo lacteus*, and pearl-spotted owlet *Glaucidium perlatum* (Hockey *et al*., 2005).

### 3.2 Data collection

In each village we visited learners in one primary school (grade 6/7; ages 12-13) and one secondary school (grade 11/12; ages 17-18). Young people were targeted because environmental education may be more effective at changing attitudes when people are exposed to concepts earlier in life (Caro *et al*., 1994), and they can also successfully change attitudes of other family members (Marchini and Macdonald, 2019). In August 2016 we administered a questionnaire (Document S1a) to a total of 283 learners at the two primary schools and two secondary schools from the two villages (Table S2). We then delivered a 20-minute presentation on the natural history of owls (slides shown in Document S3). The presentation included information on the mean number of rodents eaten by a western barn owl and a spotted eagle owl in a night (Verreaux’s eagle-owl is uncommon and pearl-spotted owls are largely insectivorous). The scholars were involved in the presentation by being asked to calculate, based on the information provided, how many rodents an individual owl could potentially eat in a year. We administered a very similar questionnaire (Document S1b) to 340 learners at the same four schools in November 2016 to assess whether perceptions had changed over the intervening three months. Seventy four of the learners that completed questionnaires in the follow up survey had not watched the presentation, and individuals belonging to this group were used as a control group. The presentation and questionnaires were conducted in English, and translated into Tshivenda and Xitsonga by a local interpreter. The questionnaire questions and responses used in the analysis are shown in Table 1. Informed consent was obtained from the principle of each school and the teachers of each class, who gave permission to participate in the study after discussing the questionnaires in detail, and answering any questions they had. The teachers were also present when the questionnaires were administered.

**Table 1.** Questions and available responses used in the questionnaire.

| Question | Response |
| --- | --- |
| 1. Would you like to have an owl nesting near your home? | Yes/no |
| 2. Would you like to have an owl nesting in the roof of your home? | Yes/no |
| 3. Would you put up an artificial nest for owls to nest in, in your yard (compound) near your home? | Yes/no |
| 4. Which of the three choices best describes your feeling towards owls? | Not like/no feeling/like |
| 5. Which of the three choices best describes your response to seeing an owl? | Not afraid/very afraid/terrified |
| 6. What do you do if an owl lands on the roof of your home? | Nothing/run away/chase the owl away/try and kill the owl |
| 7. What did you do the last time you saw an owl? | Nothing/run away/try and kill the owl |
| 8. What do you believe is going to happen if an owl lands on the roof of your home? | Open response |
| 9. What problem, if any, do rats and mice cause for you and your family? | Open response |

This research received ethical approval from the University of Venda (SMNS/14/ZOO/03/2802), and was conducted under a research permit issued by the Limpopo Department of Economic Development, Environment and Tourism (LEDET) (reference number ZA/LP/88067).

### 3.3. Data analysis

We coded data in such a way that more positive responses regarding perceptions of owls and coexisting with owls were given more positive values, in order to facilitate interpretation of the results. We tested for differences in responses to questions with binary responses (questions 1-3) by fitting Bernoulli generalised linear models to the data using the glm function in the stats package in base R version 3.6.1 (R Development Core Team, 2019). We used the conditional log-log link functions to allow for more asymmetry in the distributions. We used responses as dependant variables and stage (either before or after watching the presentation) as independent variables. To test for differences in responses to questions with multiple ranked responses (questions 4-7) we fitted ordered logistic regression models, again with responses as dependant variables and survey as independent variables, using the polr function in the package MASS (Venables and Ripley, 2002). We used Akaike’s information criterion to compare models including responses from the treatment group against null models, and models including responses from the control group against null models. Plots were created using ggplot2 (Wickham, 2016). We considered the main themes emerging in the responses to the open questions 8 and 9, and extracted representative quotes. Furthermore we categorised responses to question 8 into those that mentioned traditional cultural beliefs around owls and those that mentioned impacts of owls on controlling rodents. We compared the proportions of responses falling into these categories between respondents before watching the presentation, in the follow-up survey after the presentation was given, sub-divided between students that did see the presentation and those that did not watch the presentation. All data and R code are publicly available (Williams *et al*., 2019).

## 4. Results

Perceptions of owls were generally negative both before and after listening to the presentation (Figs 2-3). But while perceptions towards owls were still negative overall after watching the presentation, responses to questions 1-6 were less negative after watching the presentation than before watching the presentation, supporting our hypothesis (Table 2). For question 7 responses did not differ between questionnaires administered before or after watching the presentation. Models of responses of the control group to each question fitted the data no better than the null models, suggesting that any differences in the treatment group were likely to be linked to listening to the presentation.

**Fig. 2.**
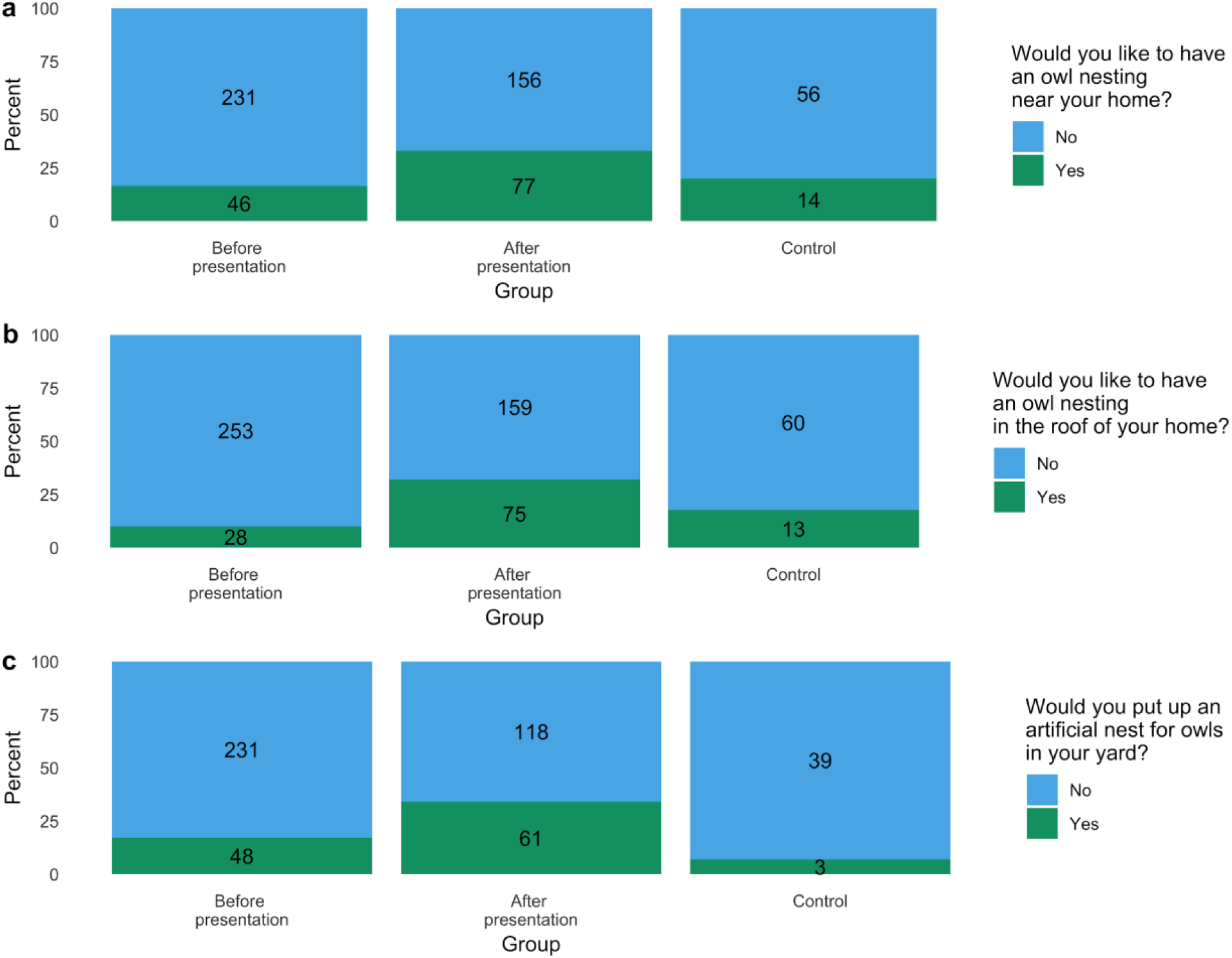
Responses to questions with binary responses (a) question 1; b) question 2; c) question 3) of three groups of learners: before watching a presentation on the natural history of owls (left); after watching the presentation (middle); and a control group that did not watch the presentation (right). Number labels show sample sizes.

**Fig. 3.**
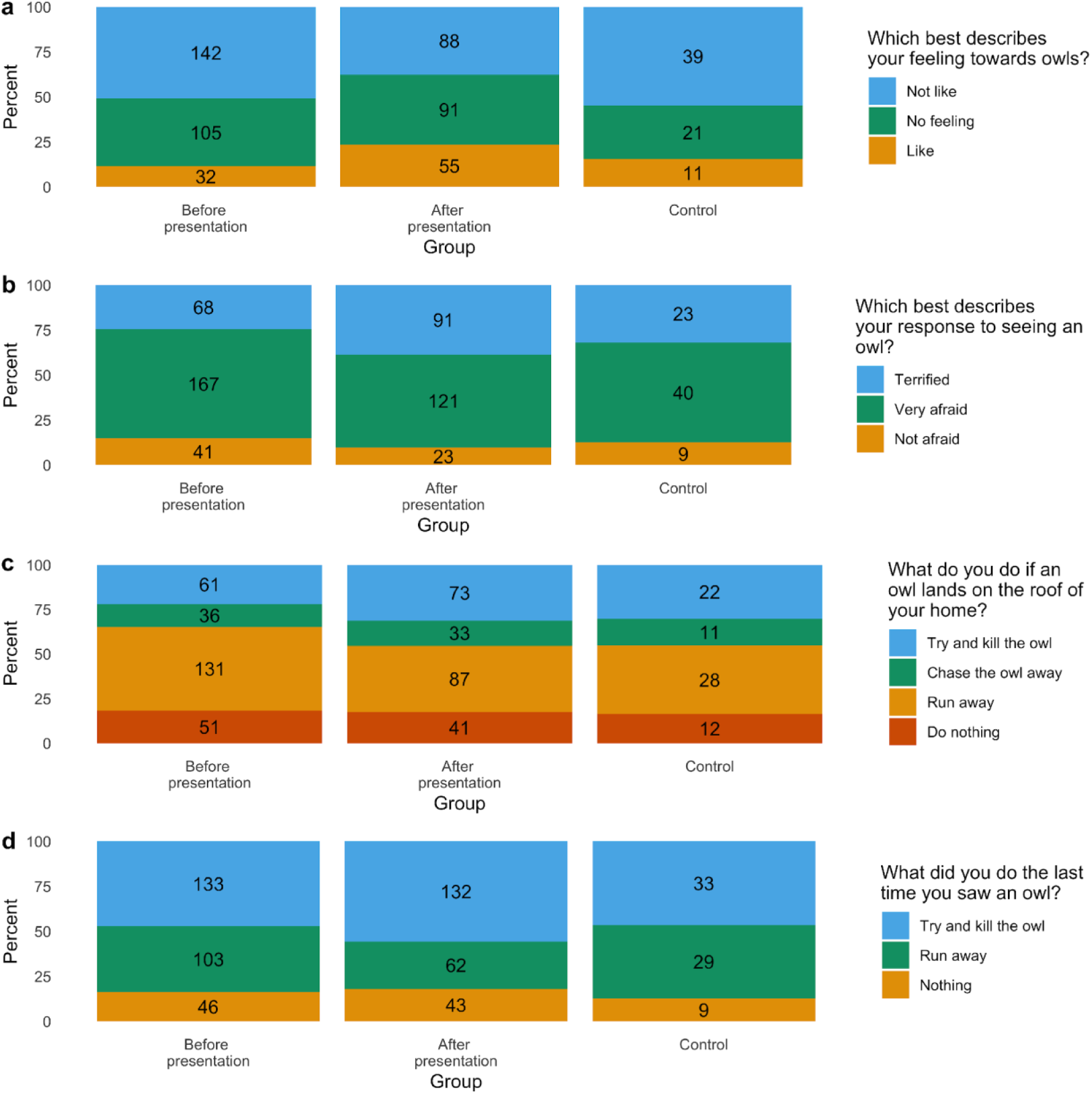
Responses to questions with more than two responses (a) question 4; b) question 5; c) question 6, d) question 7) of three groups of learners: before watching a presentation on the natural history of owls (left); after watching the presentation (middle); and a control group that did not watch the presentation (right). Number labels show sample sizes.

**Table 2.** Summary of models of responses to questions on perceptions of owls. Models with best fit, and with confidence intervals that do not overlap zero, are shown in bold.

| Question | Model type | Model name | AIC | ΔAIC | Degrees of freedom | 2.5% confidence interval | 97.5% confidence interval | Odds ratio |
| --- | --- | --- | --- | --- | --- | --- | --- | --- |
| Q1: Would you like to have an owl nesting near your home (Yes/No)? | Bernoulli GLM | **Treatment** | **548.76** | **0.00** | **2** | **0.4307009** | **1.165709** | **2.2091569** |
|  |  | Treatment null | 565.48 | 16.70 | 1 |  |  |  |
|  |  | Control | 323.13 | 0.00 | 2 | -0.4302624 | 0.7783122 | 1.2287654 |
|  |  | Control null | 321.57 | 1.60 | 1 |  |  |  |
| Q2: Would you like to have an owl nesting in the roof of your home? (Yes/No) | Bernoulli GLM | **Treatment** | **479.81** | **0.00** | **2** | **0.8808712** | **1.754406** | **3.6813819** |
|  |  | Treatment null | 517.41 | 37.60 | 1 |  |  |  |
|  |  | Control | 254.65 | 0.00 | 2 | -0.06633402 | 1.263266 | 1.8683803 |
|  |  | Control null | 255.83 | 1.20 | 1 |  |  |  |
| Q3: Would you put up an artificial nest for owls to nest in, in your yard (compound) near your home? (Yes/No) | Bernoulli GLM | **Treatment** | **489.86** | **0.00** | **2** | **0.4137402** | **1.175357** | **2.207173** |
|  |  | Treatment null | 504.66 | 14.80 | 1 |  |  |  |
|  |  | Control | 281.80 | 0.00 | 2 | -2.35006 | 0.06745143 | 0.3925334 |
|  |  | Control null | 283.07 | 1.30 | 1 |  |  |  |
| Q4: Which of the three choices best describes your feeling towards owls? (Like/No feeling/Not like) | Ordinal logistic regression | **Treatment** | **1048.78** | **0.00** | **4** | **0.3012243** | **0.9643519** | **1.881341** |
|  |  | Treatment null | 1060.96 | 12.20 | 3 |  |  |  |
|  |  | Control | 684.45 | 2.00 | 4 | -0.5778764 | 0.4458171 | 0.9419206 |
|  |  | Control null | 682.50 | 0.00 | 3 |  |  |  |
| Q5: Which of the three choices best describes your response to seeing an owl? (Not afraid/Very afraid/Terrified) | Ordinal logistic regression | **Treatment** | **963.30** | **0.00** | **4** | **0.2657433** | **0.9566639** | **1.83948** |
|  |  | Treatment null | 973.46 | 10.20 | 3 |  |  |  |
|  |  | Control | 726.65 | 0.60 | 4 | -0.7946631 | 0.222922 | 0.7507923 |
|  |  | Control null | 659.21 | 0.00 | 3 |  |  |  |
| Q6: What do you do if an owl lands on the roof of your home? (Run away/Nothing/Try and kill the owl/Chase the owl away) | Ordinal logistic regression | **Treatment** | **1331.63** | **0.00** | **5** | **0.01945016** | **0.65954542** | **1.402969** |
|  |  | Treatment null | 1333.96 | 2.30 | 4 |  |  |  |
|  |  | Control | 906.50 | 0.00 | 5 | -0.1232518 | 0.837112 | 0.837112 |
|  |  | Control null | 906.63 | 0.10 | 4 |  |  |  |
| Q7: What did you do the last time you saw an owl? (Run away/Nothing/Try and kill the owl) | Ordinal logistic regression | Treatment | 1054.50 | 0.19 | 4 | -0.1046666 | 0.5577318 | 1.253468 |
|  |  | Treatment null | 1054.31 | 0.00 | 3 |  |  |  |
|  |  | Control | 722.65 | 0.90 | 4 | -0.4380951 | 0.5392407 | 1.048204 |
|  |  | Control null | 720.69 | 0.00 | 3 |  |  |  |

The dominant theme in responses to the open questions on perceptions of owls was the involvement of owls in witchcraft, and the negative consequences that this will have for those that live in areas with owls. When asked “What do you believe is going to happen if an owl lands on the roof of your home?” typical responses include “Someone going to die”, “That someone is about to bewitch me”, “I believe that it is sent by witches”, or “Nothing happy, I will chase it away”. Some respondents also expressed more utilitarian views such as “It will help me killing rats” or “I will just try to kill it because it make a problem - noise”. In contrast, when asked “What problem, if any, do rats and mice cause for you and your family?”, respondents were less likely to share supernatural beliefs, again focussing on utilitarian impacts such as “Eat food, clothes, baby, door, everything”, “[Make me] sick”, or “Bring owl and snakes at home”. After watching the presentation answers given in response to the question “What do you believe is going to happen if an owl lands on the roof of your home?” shifted. Prior to the presentation, 52.6% of responses (n=228) expressed that the owl was sent by witches and would bring bad omens upon their household, and 13.2% of responses suggested that the owl would kill rodents. Other responses such as utilitarian concerns and worries about owls making noises comprised the remainder of responses. Following the environmental education programme, 20.7% of responses given by presentation attendees (n = 92) pertained to witchcraft and ill omens while 27.1% of responses focussed on predation of mice and rats by owls.

## 5. Discussion

Perceptions of owls were less negative after watching the presentation than before watching the presentation. Despite this shift, perceptions of owls still remained negative overall, which was linked to their associations with witchcraft, although the prevalence of responses relating to a negative association between owls and witchcraft appeared to be lower among respondents that had seen the presentation than those that had not. This is not surprising given how strongly-held beliefs in the supernatural tend to be (Dickman and Hazzah, 2016) and the low intensity with which the education programme was implemented. But these findings nevertheless demonstrate that even modest educational programmes that involve the delivery of only a single presentation can reduce negative perceptions of culturally stigmatised wildlife. A more intensive environmental education programme involving more sessions would be likely to improve perceptions of wildlife further (Kruse and Card, 2004), and although these can be costly and time-intensive to implement (Leisher *et al*., 2012), a longer term approach is recommended for species with strong negative cultural associations.

The improved attitudes of participants towards owls fits well with previous findings that participation in an environmental education scheme involving talks and activities about lemurs (superfamily Lemuroidea) in Madagascar during a single day was sufficient to improve knowledge and attitudes of school children in relation to these species (Rakotomamonjy *et al*., 2015). Similarly, an environmental education programme that centred on screening three 20-minute educational films on the threats posed to mountain gorillas *Gorilla beringei beringei* and chimpanzees *Pan troglodytes schweinfurthii* for school children in Uganda was able improve attitudes towards great apes and knowledge of conservation actions (Leeds *et al*., 2017). In these communities, however, the focal species are not strongly linked with negative supernatural superstitions (Leeds *et al*., 2017; Rakotomamonjy *et al*., 2015).

Indigenous knowledge systems play an important role in conservation and environmental education, and their integration can be advantageous, especially if traditional beliefs benefit wildlife and communities (Maila and Loubser, 2003; Risiro *et al*., 2013). Perceptions stemming from traditional beliefs or superstitions can also pose challenges to human-wildlife coexistence, which manifest in multiple forms such as right of passage ritual killings of animals (Hazzah *et al*., 2009), trade in endangered species for traditional medicine (Williams *et al*., 2013), and persecution of animals associated with bad omens or demonstrating taboo behaviours (Forth, 2007). In our study area and in other examples, negative perceptions of owls can result in damage to ecosystems and missed opportunities for farming communities to benefit from ERBM programmes (Mikkola and Mikkola, 1997; Ogada and Kibuthu, 2008). Environmental education programmes need to identify barriers that limit stakeholder engagement and carefully consider strategies to overcome these (Offord-Woolley *et al*., 2016). A culturally sensitive approach is required when addressing concepts surrounding witchcraft (Ashforth, 1996; Cumes, 2004). Outsiders may fail to grasp the dynamic and modern applications of witchcraft and cause offence by ignoring or misrepresenting the concept’s sensitive and secretive characteristics (Geschiere, 1997). If possible, conservation organisations should include community members in the design and implementation of environmental education programmes to guarantee that locally specific cultural perspectives and priorities are incorporated (Jacobson *et al*., 2015; Offord-Woolley *et al*., 2016). For owls and many other species, a wide variety of contradictory beliefs are associated with the species globally (Enriquez and Mikkola, 1997; Forth, 2007; Glickman, 1995), demonstrating the importance of ascertaining a local perspective in conservation initiatives. Hence our environmental education programme partnered with a local translator and focussed on ecology and ecosystem services of owls, rather than attempting to dissuade participants from beliefs in witchcraft. It is also important to allow time for people to readjust to new information and perceptions of a historically disliked species (Linnell *et al*., 2003), especially when perceptions have existed in communicative memory and cultural memory for long periods (Assmann and Czaplicka, 1995). An interesting extension of this study would be to assess the programme’s longer-term effectiveness on changing perceptions. Furthermore, delivery of the education programme by teaching staff at the schools may prove more effective than delivery by an external scientist, as this could signal that the message was socially accepted (Marchini and Macdonald, 2019).

In addition to superstitious views of owls, respondents tended to frame positive and negative views pertaining to owls and rodents in utilitarian terms. This is not surprising, as lower income communities have a more pressing urgency to fulfil basic needs than higher income groups, and are consequently more likely to consider animals from a utilitarian perspective (Infield, 1988). Wildlife perceived to be devoid of a useful purpose is seldom considered worthy of preservation by lower income communities (Griffiths, 2017; Williams, 2017). In South Africa, financial inequality, poverty, and economic marginalisation of certain groups from viewing wildlife has resulted in widespread attitudes that wildlife has little value (Griffiths, 2017). Within South Africa, beliefs in witchcraft and supernatural powers are more commonplace in impoverished areas, specifically parts of Limpopo and Eastern Cape provinces (Ashforth, 1996; Kohnert, 2003; Niehaus *et al*., 2001). High levels of poverty and prevalent beliefs in witchcraft amplify owl vulnerability in Limpopo province. Environmental education schemes focussing on animals associated with witchcraft should assign these species with positive, sustainable, and accessible utilitarian values, such as those owls generate in the context of EBRM, to promote species conservation.

Our findings also suggest that in addition to enhancing perceptions of wildlife, environmental education could be also an incredibly useful tool to increase participation in community programmes such as EBRM initiatives. Respondents that watched the presentation were more likely to say they would be willing to have an owl box installed in their yard, which could help reduce rodent densities in fields by increasing owl populations in agricultural areas (Paz *et al*., 2013). Although the animal welfare implications of using indigenous predators to control pests rather than relying on poison have recently been questioned (Allen *et al*., 2019), there is little doubt that using ecosystem services provided by natural predators would be more ecologically sound and more sustainable than chemical rodenticides for community members farming in rural agro-ecosystems (Singleton *et al*., 1999).

We note, however, that while we observed increased theoretical willingness to participate in a future EBRM programme involving erecting owl nesting boxes, further studies are required to assess whether this would translate into actual increased participation after the launch of such a scheme, as this is not always the case (Waylen *et al*., 2009; Young *et al*., 2013). Engaging students in constructing, erecting, and potentially monitoring owl boxes provides a constructive extension to a low intensity environmental education programme, and fortifies concepts presented in the classroom through empowering actions. An environmental education programme conducted in a semi-urban area in Gauteng province, South Africa, engaged students in a similar multi-pronged approach and following the experience, both the students involved and their families replaced superstitious beliefs about owls with more positive perspectives (Meyer, 2008). Another limitation of our study was the relatively small sample size, so follow-up studies would

In conclusion, our findings indicate that even a low intensity environmental education programme can improve young people’s perceptions of a species associated with witchcraft, and their propensity to undertake positive environmental actions. In Africa, beliefs in witchcraft and the supernatural have evolved in response to shifting politics and modernisation (McEwan, 2008; Niehaus *et al*., 2001). Through culturally sensitive and locally inclusive environmental education, negative perceptions of animals affiliated with witchcraft can also evolve to benefit communities, farmers, and wildlife.

## 7. Acknowledgements

We would like to thank the learners that responded to our questionnaires, and the teachers from Thomas Ntshavheni Primary School and Tshinavhe Secondary School, in Vyeboom, and Anderson Memorial Primary School and Ndengeza Senior Secondary School, in Ndengeza, for their support. STW was supported by a postdoctoral grant from the University of Venda. KSW was funded by the Durham University COFUND research fellowship. PJT acknowledges the financial support of the National Research Foundation and the Department of Science and Technology through the South African Research Chair on Biodiversity Value and Change (Grant No. 87311) hosted by the University of Venda and co-hosted by the Centre for Invasion Biology at Stellenbosch University. NC was supported by a postdoctoral grant administered by the National Research Foundation under the South African Research Chair on Biodiversity Value and Change at the University of Venda. LHS was supported by the National Research Foundation of South Africa (Grant No. 107099) and the University of Venda. This research was partly funded by a European Union 9th European Development Fund grant from the African Caribbean and Pacific Science and Technology Programme (StopRats: FED/2013/330-223) and by an African Union grant (EcoRodMan: AURGII/1/006/2016). The funders had no role in study design, data collection and analysis, decision to publish, or preparation of the manuscript. The authors declare that no competing interests exist.

## 8. Supplementary information

Note that all supplementary information is available in a Figshare repository (https://figshare.com/s/711295efc8d5f96b21f3; doi: p10.6084/m9.figshare.9734984). This repository will be made publicly accessible upon acceptance of the manuscript. In addition to Document S1 and Table S2 all data and code will also be accessible in the repository.

Document S1. Questionnaire administered to learners before and after listening to a presentation on the natural history of owls.

Table S2. Number of respondents at each school that completed the questionnaire before and after listening to the presentation.

## 9. Statements

### Data availability

Data available from the Figshare digital repository Williams et al. (2019) (https://figshare.com/s/711295efc8d5f96b21f3; doi: p10.6084/m9.figshare.9734984). This will be made publicly accessible upon acceptance of the manuscript.

### Authors’ contributions

SWE, LHS, PJT, and SRB conceived the ideas and designed methodology; SWE collected the data; STW analysed the data; STW, KSW, and NLC led the writing of the manuscript. All authors contributed critically to the drafts and gave final approval for publication.

